# Unique genomic features and deeply-conserved functions of long non-coding RNAs in the Cancer LncRNA Census (CLC)

**DOI:** 10.1101/152769

**Authors:** Joana Carlevaro-Fita, Andrés Lanzós, Lars Feuerbach, Chen Hong, David Mas-Ponte, Jakob Skou Pedersen, Rory Johnson, On behalf of the PCAWG Drivers and Functional Interpretation Group and the ICGC/TCGA Pan-Cancer Analysis of Whole Genomes Network

## Abstract

Long non-coding RNAs (lncRNAs) that drive tumorigenesis are a growing focus of cancer genomics studies. To facilitate further discovery, we have created the “Cancer LncRNA Census” (CLC), a manually-curated and strictly-defined compilation of lncRNAs with causative roles in cancer. CLC has two principle applications: first, as a resource for training and benchmarking *de novo* identification methods; and second, as a dataset for studying the fundamental properties of these genes.

CLC Version 1 comprises 122 lncRNAs implicated in 29 distinct cancers. LncRNAs are included based on functional or genetic evidence for causative roles in cancer progression. All belong to the GENCODE reference annotation, to enable integration across projects and datasets. For each entry, the evidence type, biological activity (oncogene or tumour suppressor), source reference and cancer type are recorded. Supporting its usefulness, CLC genes are significantly enriched amongst *de novo* predicted driver genes from PCAWG. CLC genes are distinguished from other lncRNAs by a series of features consistent with biological function, including gene length, high expression and sequence conservation of both exons and promoters. We identify a trend for CLC genes to be co-localised with known protein-coding cancer genes along the human genome. Finally, by integrating data from transposon-mutagenesis functional screens, we show that mouse orthologues of CLC genes tend also to be cancer genes.

Thus CLC represents a valuable resource for research into long non-coding RNAs in cancer. Their evolutionary and genomic properties have implications for understanding disease mechanisms and point to conserved functions across ~80 million years of evolution.

## Introduction

Tumorigenesis is driven by a series of genetic mutations that promote cancer phenotypes and consequently experience positive selection (Yates & Campbell 2012). The systematic discovery of such driver mutations, and the genes whose functions they alter, has been made possible by tumour genome sequencing. By collecting the entirety of such genes for every cancer type, we aim to develop a comprehensive view of underlying processes and pathways, and thereby formulate effective, targeted therapeutic strategies.

The cast of genetic elements implicated in tumorigenesis has recently grown as diverse new classes of non-coding RNAs and regulatory features have been discovered. These include the long non-coding RNAs (lncRNAs), of which tens of thousands have been catalogued (Guttman et al. 2009; Jia et al. 2010; Cabili et al. 2011; Derrien et al. 2012). LncRNAs are >200 nt long transcripts with no protein-coding capacity. Their evolutionary conservation and regulated expression, combined with a number of well-characterised examples, have together led to the view that lncRNAs are *bona fide* functional genes (Grote et al. 2013; Sauvageau et al. 2013; Ulitsky & Bartel 2013; Liu et al. 2017). Current thinking holds that lncRNAs function by forming complexes with proteins and RNA both inside and outside the nucleus (Guttman & Rinn 2012; Johnson & Guigó 2014).

LncRNAs have been shown to play important roles in various cancers. For example, *MALAT1*, a potent oncogene across numerous cancers, is restricted to the nucleus and plays a housekeeping role in splicing (Gutschner & Diederichs 2012; Engreitz et al. 2014). *MALAT1* is overexpressed in a variety of cancer types, and its knockdown potently reduces not only proliferation but also metastasis *in vivo* (Gutschner et al. 2013). *MALAT1* gene is subjected to elevated mutational rates in human tumours, although it has not yet been established whether these mutations drive tumorigenesis (Lanzós et al. 2017) (PCAWG Consortium, Manuscript in Preparation). On the other hand, lncRNAs may also function as tumour suppressors. *LincRNA-p21* acts as a downstream effector of p53 regulation through recruitment of the repressor hnRNP-K (Huarte et al. 2010). These and other examples of lncRNAs linked to cancer, raise the question of how many more remain to be found amongst the ~99% of lncRNAs that are presently uncharacterised (Derrien et al. 2012; Quek et al. 2015; Iyer et al. 2015).

Recent tumour genome sequencing studies, in step with advanced bioinformatic driver-gene prediction methods, have yielded hundreds of new candidate protein-coding driver genes (Tamborero et al. 2013). For economic reasons, these studies initially restricted their attention to “exomes” or the ~2% of the genome covering protein-coding exons (Chang et al. 2013). Unfortunately such a strategy ignores mutations in the remaining ~98% of genomic sequence, home to the majority of lncRNAs (Gutschner & Diederichs 2012; Derrien et al. 2012). Driver gene identification methods rely on statistical models that make a series of assumptions about and simplifications of complex tumour mutation patterns (Lawrence et al. 2014). It is critical to test the performance of such methods using true-positive lists of known cancer driver genes. For protein-coding genes, this role has been fulfilled by the Cancer Gene Census (CGC) (Futreal et al. 2004), which is collected and regularly updated by manual annotators. Comparison of driver predictions to CGC genes facilitates further method refinement and comparison between methods (Sjoblom et al. 2006; Redon et al. 2006; Mularoni et al. 2016; Tokheim et al. 2016).

In addition to its benchmarking role, the CGC resource has also been useful in identifying unique biological features of cancer genes. For example, CGC genes tend to be more conserved and longer. Furthermore, they are enriched for genes with transcription regulator activity and nucleic acid binding functions (Furney et al. 2006; Furney et al. 2008).

Until very recently, efforts to discover cancer lncRNAs have depended on classical functional genomics approaches of differential expression using microarrays or RNA sequencing (Huarte et al. 2010; Iyer et al. 2015). While valuable, differential expression *per se* is not direct evidence for causative roles in tumour evolution. To more directly identify lncRNAs that drive cancer progression, a number of methods, including several within the PCAWG Network (PCAWG Consortium, Manuscript in Preparation), have recently been developed to search for signals of positive selection using mutation maps of tumour genomes. OncodriveFML utilises nucleotide-level functional impact scores inferred from predicted changes in RNA secondary structure (Sabarinathan et al. 2013) together with an empirical significance estimate, to identify lncRNAs with an excess of high-impact mutations (Mularoni et al. 2016). Another method, ExInAtor, identifies candidates with elevated mutational load, using trinucleotide-adjusted local background (Lanzós et al. 2017). A clear impediment in both cases has been the lack of true-positive set of known lncRNA driver genes, analogous to CGC. Although there do exist databases of cancer lncRNAs, notably LncRNADisease (Chen et al. 2013) and Lnc2Cancer (Ning et al. 2016), they mix unfiltered data from numerous sources, resulting in inconsistent criteria for inclusion (including expression changes), and inconsistent gene identifiers.

To facilitate the future discovery of cancer lncRNAs, and gain insights into their biology, we have compiled a highly-curated set of cases with roles in cancer processes. Here we present the *Cancer LncRNA Census* (CLC), the first compendium of lncRNAs with direct functional or genetic evidence for cancer roles. We demonstrate the utility of CLC in assessing the performance of driver lncRNA predictions. Through analysis of this geneset, we demonstrate that cancer lncRNAs have a unique series of features that may in future be used to assist *de novo* predictions. Finally, we show that CLC genes have conserved cancer roles across the approximately 80 million years of evolution separating humans and rodents.

## Results

### Definition of cancer related lncRNAs

As part of recent efforts to identify driver lncRNAs by the Drivers and Functional Interpretation Group (PCAWG-2-5-9-14) within the ICGC/TCGA Pan-Cancer Analysis of Whole Genomes Network (henceforth PCAWG), we discovered the need for a high-confidence reference set of cancer-related lncRNA genes, which we henceforth refer to as “cancer lncRNAs”. We here present Version 1 of the *Cancer LncRNA Census* (CLC).

Cancer lncRNAs were identified from the literature using defined and consistent criteria, being direct experimental or genetic evidence for roles in cancer progression or phenotypes (see Materials and Methods). Alterations in expression alone were not considered sufficient evidence. Importantly, only lncRNAs with GENCODE identifiers were included. For every cancer lncRNA, one or more associated cancer types were collected.

Attesting to the value of this approach, we identified several cases in semi-automatically annotated cancer lncRNA databases of lncRNAs that were misassigned GENCODE identifiers, usually with an overlapping protein-coding gene (Chen et al. 2013). We also excluded a number of published lncRNAs for which we could not find evidence to meet our criteria, for example *CONCR*, *SRA1* and *KCNQ1OT1* (Marchese et al. 2016; Lanz et al. 1999; Higashimoto et al. 2006).

Version 1 of CLC contains 122 lncRNA genes, however, eight of them are annotated as pseudogenes rather than lncRNAs by GENCODE. The remaining 114 CLC genes correspond to 0.72% of a total of 15,941 lncRNA gene loci annotated in GENCODE v24 (Derrien et al. 2012; Harrow et al. 2012) (Figure 1). For comparison, the Cancer Gene Census (CGC) (COSMIC v78, downloaded Oct, 3, 2016) lists 561 or 2.8% of protein-coding genes (Futreal et al. 2004). The entire remaining set of 15,827 lncRNA loci is henceforth referred to as “nonCLC” (Figure 1). The full CLC dataset is found in Supplementary Table 1.

**Table 1:**
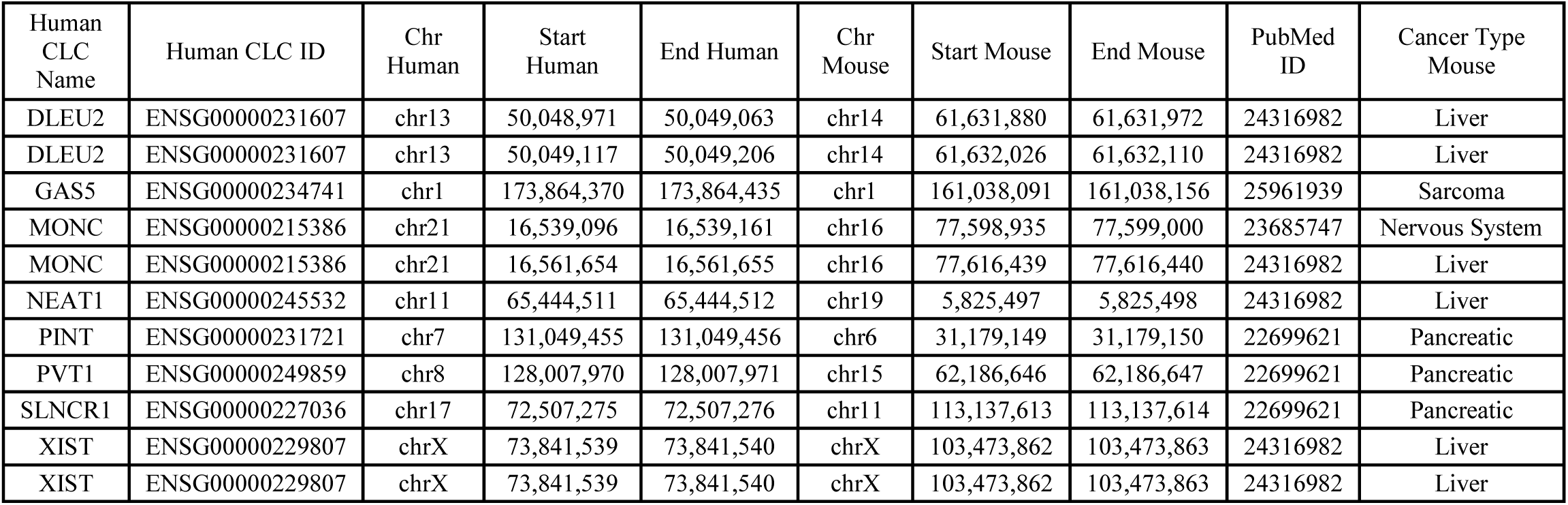
List of intergenic CIS human (GRCh38) / mouse (GRCm38) gene pairs.

| Human<br>CLC<br>Name | Human CLC ID | Chr<br>Human | Start<br>Human | End Human | Chr<br>Mouse | Start Mouse | End Mouse | PubMed<br>ID | Cancer Type<br>Mouse |
| --- | --- | --- | --- | --- | --- | --- | --- | --- | --- |
| DLEU2 | ENSG00000231607 | chr13 | 50,048,971 | 50,049,063 | chr14 | 61,631,880 | 61,631,972 | 24316982 | Liver |
| DLEU2 | ENSG00000231607 | chr13 | 50,049,117 | 50,049,206 | chr14 | 61,632,026 | 61,632,110 | 24316982 | Liver |
| GAS5 | ENSG00000234741 | chr1 | 173,864,370 | 173,864,435 | chr1 | 161,038,091 | 161,038,156 | 25961939 | Sarcoma |
| MONC | ENSG00000215386 | chr21 | 16,539,096 | 16,539,161 | chr16 | 77,598,935 | 77,599,000 | 23685747 | Nervous System |
| MONC | ENSG00000215386 | chr21 | 16,561,654 | 16,561,655 | chr16 | 77,616,439 | 77,616,440 | 24316982 | Liver |
| NEAT1 | ENSG00000245532 | chr11 | 65,444,511 | 65,444,512 | chr19 | 5,825,497 | 5,825,498 | 24316982 | Liver |
| PINT | ENSG00000231721 | chr7 | 131,049,455 | 131,049,456 | chr6 | 31,179,149 | 31,179,150 | 22699621 | Pancreatic |
| PVT1 | ENSG00000249859 | chr8 | 128,007,970 | 128,007,971 | chr15 | 62,186,646 | 62,186,647 | 22699621 | Pancreatic |
| SLNCR1 | ENSG00000227036 | chr17 | 72,507,275 | 72,507,276 | chr11 | 113,137,613 | 113,137,614 | 22699621 | Pancreatic |
| XIST | ENSG00000229807 | chrX | 73,841,539 | 73,841,540 | chrX | 103,473,862 | 103,473,863 | 24316982 | Liver |
| XIST | ENSG00000229807 | chrX | 73,841,539 | 73,841,540 | chrX | 103,473,862 | 103,473,863 | 24316982 | Liver |

**Figure 1:**
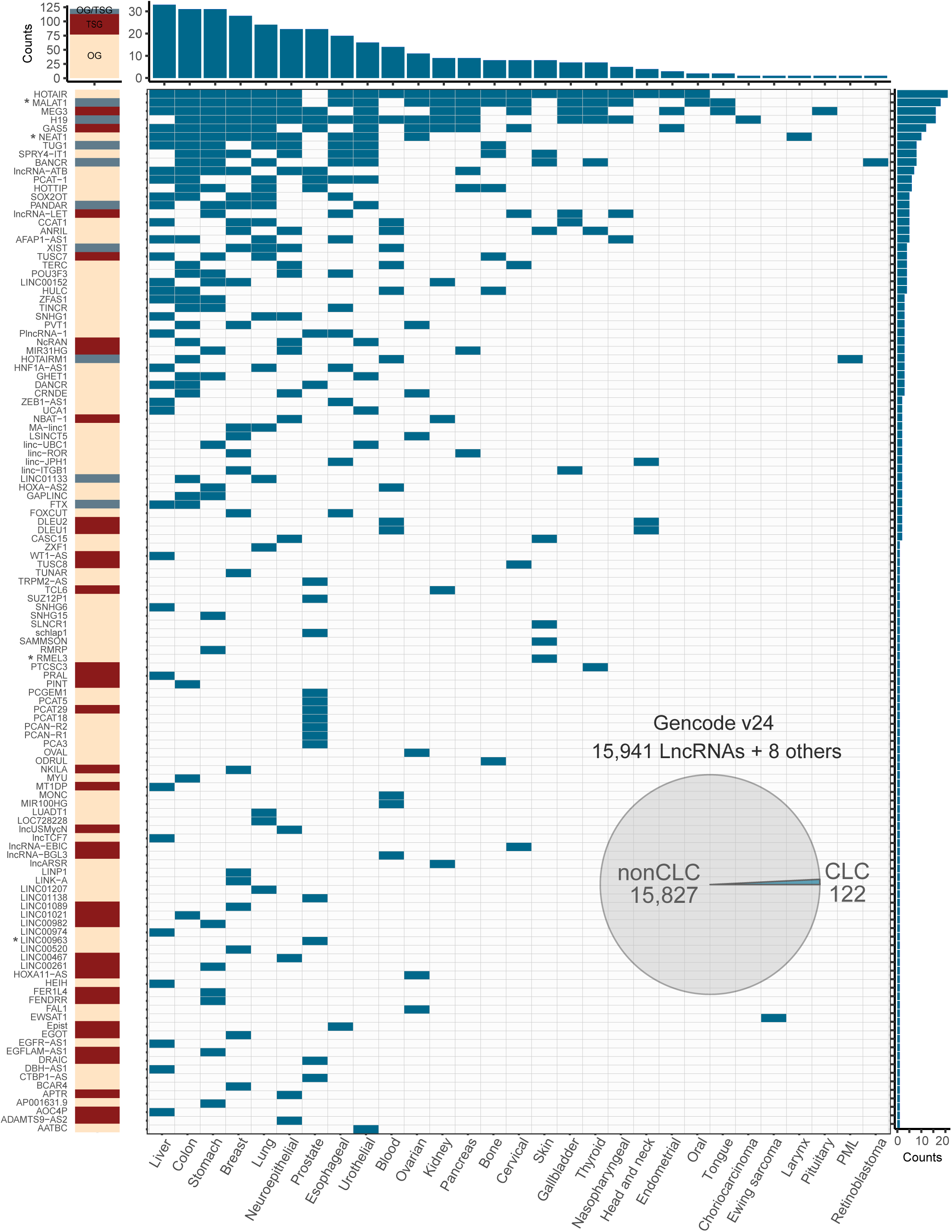
Overview of the Cancer LncRNA Census. Rows represent the 122 CLC genes, columns represent 29 cancer types. Asterisk next to gene names indicate that they are predicted as drivers by PCAWG, based either on gene or promoter evidence (see Supplementary Table 1).. Blue cells indicate evidence for the involvement of a given lncRNA in that cancer type. Left column indicates functional classification: tumour suppressor (TSG), oncogene (OG) or both (OG/TSG). Above and to the right, barplots indicate the count totals of each column / row. The piechart shows the fraction that CLC within GENCODE v24 lncRNAs. Note that 8 CLC genes are classified as “pseudogenes” by GENCODE. “nonCLC” refers to all other GENCODE-annotated lncRNAs, which are used as background in comparative analyses.

The cancer classification terminology used amongst the source literature for CLC was not uniform. Therefore, using the International Classification of Diseases for Oncology (World Health Organization 2013), we reassigned the cancer types described in the original research articles to a reduced set of 29 (Figure 1 and Supplementary Figure 1).

Altogether, CLC contains 333 unique lncRNA-cancer type relationships. Out of 122 genes, 77 (63.1%) were shown to function as oncogenes, 36 (29.5%) as tumour suppressors, and 9 (7.4%) with evidence for both activities (Figure 1 and Supplementary Figure 1).

The most prolific lncRNAs, with >=16 recorded cancer types, are *HOTAIR*, *MALAT1*, *MEG3* and *H19* (Figure 1 and Supplementary Figure 1). It is not clear whether this reflects their unique pan-cancer functionality, or is simply a result of their being amongst the most early-discovered and widely-studied lncRNAs.

*In vitro* experiments were the most frequent evidence source, usually consisting of RNAi-mediated knockdown in cultured cell lines, coupled to phenotypic assays such as proliferation or migration (Supplementary Figure 1). Far fewer have been studied *in vivo*, or have cancer-associated somatic or germline mutations. 19 lncRNAs had 3 or more independent evidence sources (Supplementary Figure 1).

### CLC and other databases

There are a number of relevant lncRNA databases presently available: the *Lnc2Cancer* database (n= 654) (Ning et al. 2016), the *LncRNADisease* Database (n=121) (Chen et al. 2013), *lncRNAdb* (n=191) (Quek et al. 2015) and the “Cancer Related LncRNAs” set we recently produced (n=45) (Lanzós et al. 2017). CLC covers between 17% and 31% of these databases (*Lnc2Cancer* and *LncRNADisease* respectively) but none of these resources contain the complete list of genes presented here (Figure 2A). We sought to use recent unbiased proliferation screen data to independently compare cancer lncRNA databases (Zhu et al. 2016; Liu et al. 2017). Using only GENCODE-annotated genes, CLC is the resource that overall has the highest fraction of independently-identified proliferation lncRNAs, although the sparse nature of the data means that this conclusion is not definitive (Figure 2B).

**Figure 2:**
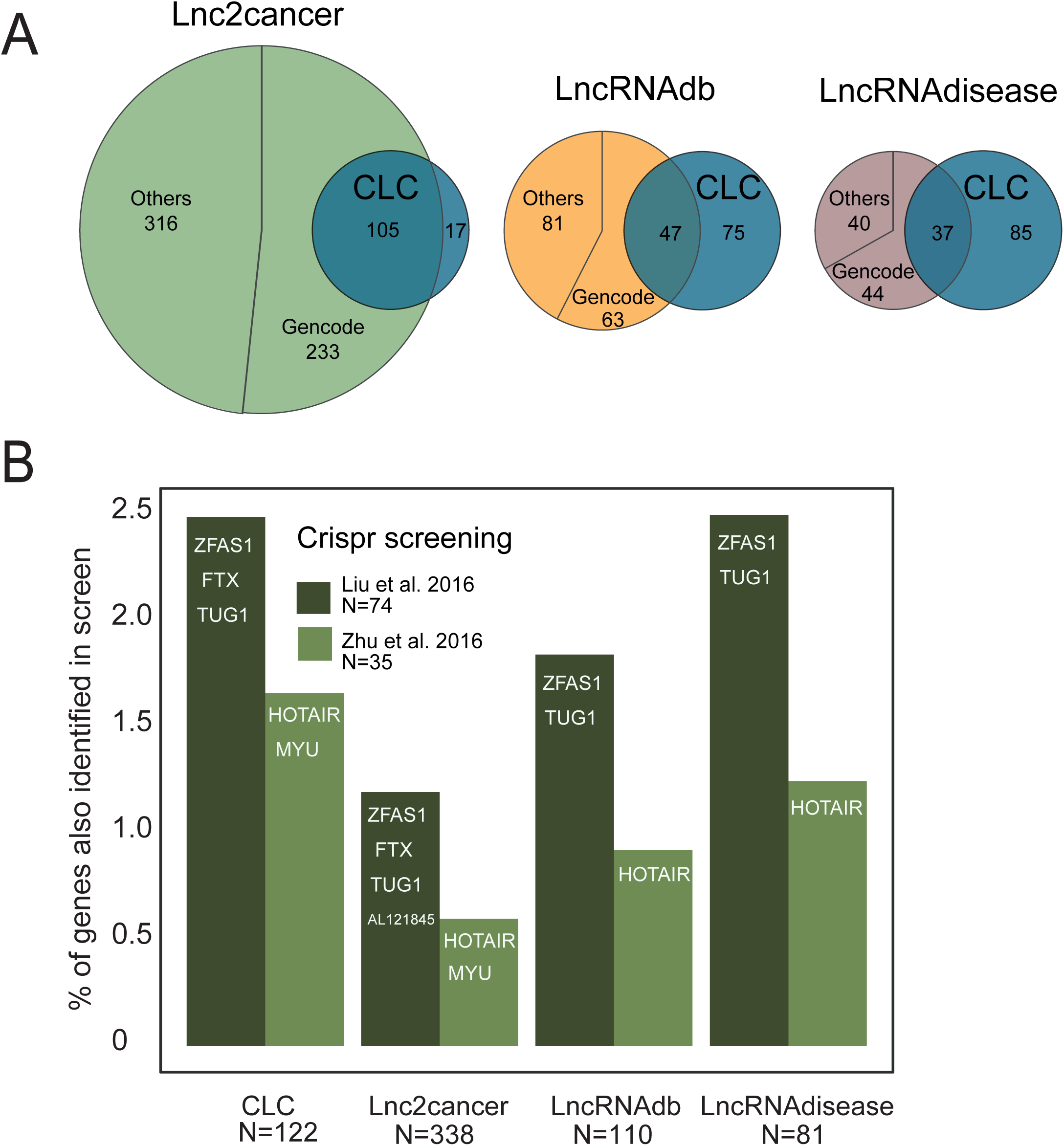
Intersection of CLC with public databases. (A) Proportional Venn diagrams displaying the overlap between CLC set and the three indicated databases. Shown are the total number of unique human lncRNAs contained in each intersection (note that for LncRNADisease, numbers refer only to cancer-related genes). Databases are divided into genes that belong to GENCODE v24 annotation and others. (B) Barplot shows the percent of GENCODE v24 lncRNAs of each database that is present in the final list of cancer lncRNA candidates of two CRISPR cancer screenings (Liu et al. 2016 and Zhu et al. 2016). N represents the number of GENCODE v24 lncRNAs that could be used for the analysis. Names of the genes that overlap between the databases and the screenings are shown in each bar.

### CLC for benchmarking lncRNA driver prediction methods

One of the primary motivations for CLC is to develop a true positive set for benchmarking and comparing methods for identifying driver lncRNAs. In the domain of protein-coding driver gene predictions, the Cancer Gene Census (CGC) has become such a “gold standard” training set (Futreal et al. 2004). Typically, the predicted driver genes belonging to CGC are judged to be true positives, and the fraction of these amongst predictions is used to estimate the Positive Predictive Value (PPV), or precision. This measure can be calculated for increasing cutoff levels, to assess the optimal cutoff.

First, we used CLC to examine the performance of the lncRNA driver predictor ExInAtor (Lanzós et al. 2017) in recalling CLC genes using PCAWG tumour mutation data (PCAWG Consortium, Manuscript In Preparation). A total of 2,687 GENCODE lncRNAs were tested here, of which 82 (3.1%) belong to CLC. Driver predictions on several cancers at the standard False Discovery Rate (“*q*-value”) cutoff of 0.1 are shown for selected cancers in Figure 3A. That panel shows the CLC-defined precision (*y*-axis) as a function of predicted driver genes ranked by *q*-value (*x*-axis). We observe rather heterogeneous performance across cancer cohorts. This may reflect a combination of intrinsic biological differences and differences in cohort sizes, which differs widely between the datasets shown. For the merged pan-cancer dataset, ExInAtor predicted three CLC genes amongst its top ten candidates (*q*-value < 0.1), a rate far in excess of the background expectation (“Baseline”, being the fraction all lncRNAs being in CLC). Similar enrichments are observed for other cancer types. These results support both the predictive value of ExInAtor, and the usefulness of CLC in assessing lncRNA driver predictors. Comprehensive CLC-based assessments of lncRNA driver discovery, across all methods and tumour cohorts in PCAWG, may be found in the main PCAWG driver prediction publication (PCAWG Consortium, Manuscript In Preparation).

**Figure 3:**
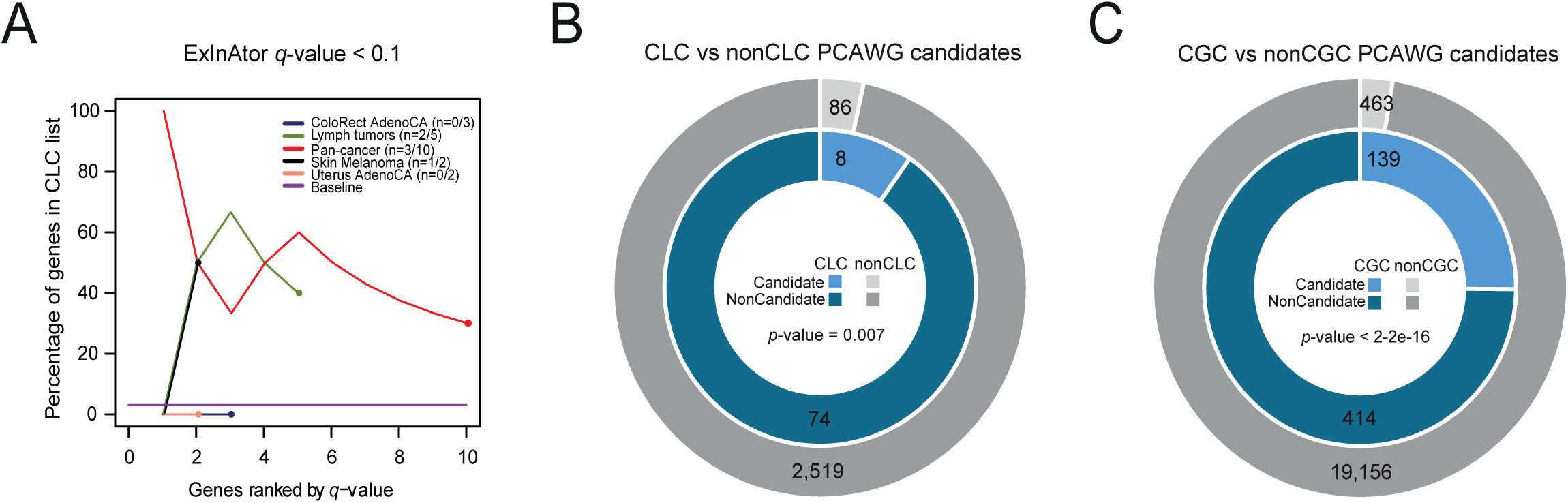
CLC as benchmark for cancer driver predictions. (A) CLC benchmarking of ExInAtor driver lncRNA predictions using PCAWG whole genome tumours at *q*-value (false discovery rate) cutoff of 0.1. Genes sorted increasingly by *q*-value are ranked on *x* axis. Percentage of CLC genes amongst cumulative set of predicted candidates at each step of the ranking (precision), are shown on the *y* axis. Black line shows the baseline, being the percentage of CLC genes in the whole list of genes tested. Coloured dots represent the number of candidates predicted under the *q*-value cutoff of 0.1. “n” in the legend shows the number of CLC and total candidates for each cancer type. (B) Rate of driver gene predictions amongst CLC and nonCLC genesets (*q*-value cutoff of 0.1) by all the individual methods and the combined list of drivers developed in PCAWG. *p*-value is calculated using Fisher’s exact test for the difference between CLC and nonCLC genesets. (C) Rate of driver gene predictions amongst CGC and nonCGC genesets (*q*-value cutoff of 0.1) by all the individual methods and the combined list of drivers developed in PCAWG. *p*-value is calculated using Fisher’s exact test for the difference between CGC and nonCGC genesets.

Finally, we assessed the precision (i.e. positive predictive value) of PCAWG lncRNA and protein-coding driver predictions across all cancers and all prediction methods (PCAWG Consortium, Manuscript In Preparation). Using the same *q*-value cutoff of 0.1, we found that across all cancer types and methods, a total of 8 (8.5%) of lncRNA predictions belong to CLC (Figure 3B), while a total of 139 (23.1%) of protein-coding predictions belong to CGC (Figure 3C). In terms of sensitivity, 9.8% and 25.1% of CLC and CGC genes are predicted as candidates, respectively. Despite the lower detection of CLC genes in comparison to CGC genes, both sensitivity rates significantly exceed the prediction rate of nonCLC and nonCGC genes (P=0.007 and P<0.001 Fisher’s exact tests, respectively), again highlighting the usefulness of the CLC geneset (Figure 3C).

### CLC genes are distinguished by function- and disease-related features

We recently found evidence, using a smaller set of Cancer Related LncRNAs (CRLs), that cancer lncRNAs are distinguished by various genomic and expression features indicative of biological function (Lanzós et al. 2017). We here extended these findings using a large series of potential gene features, to search for those features distinguishing CLC from nonCLC lncRNAs (Figure 4A).

**Figure 4:**
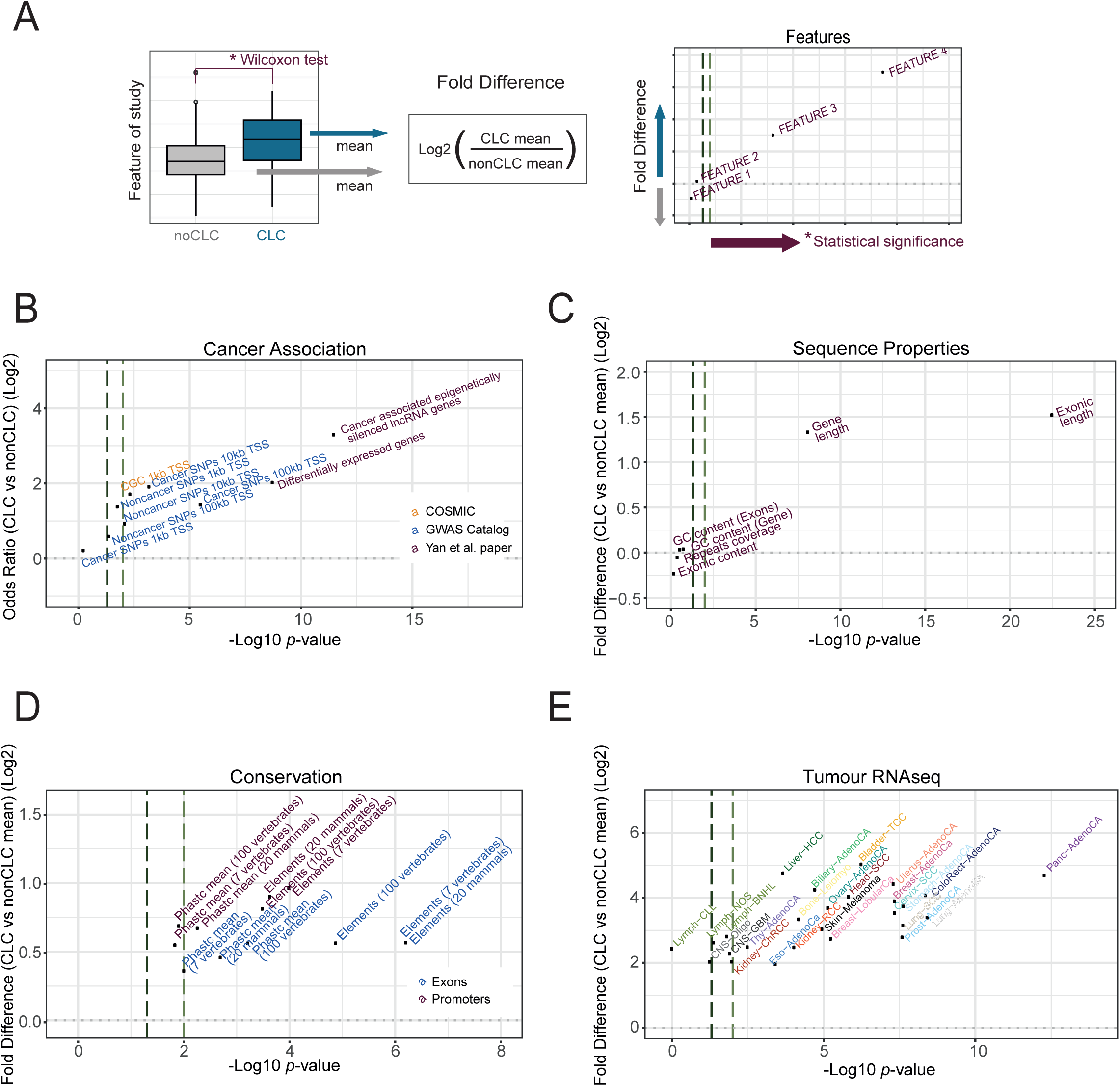
Distinguishing features of CLC genes. (A) Panel showing a hypothetic feature analysis example to illustrate the content of the following figures. All panels in figure 4 display features (dots), plotted by their log fold difference (Odds Ratio in case of panel “B”) between CLC / nonCLC genesets (*y*-axis) and statistical significance (*x*-axis). In all plots dark and light green dashed lines indicate 0.05 and 0.01 significance thresholds, respectively. (B) Cancer and noncancer disease related data from indicated sources: *y*-axis shows the log2 of the Odds Ratio obtained by comparing CLC to nonCLC by Fisher’s exact test; *x-*axis displays the estimated *p*-value from the same test. “CGC 1 kb TSS” refers to the fraction of genes that have a nearby known CGC cancer protein-coding gene. This is explored in more detail in the next Figure. “Noncancer SNPs” refers to GWAS SNPs associated with diseases/traits other than cancer (C) Sequence and gene properties: *y*-axis shows the log2 fold-difference of CLC / nonCLC means; *x*-axis represents the *p*-value obtained. (D) Evolutionary conservation: “Phastc mean” indicates average base-level PhastCons score; “Elements” indicates percent coverage by PhastCons conserved elements (see Materials and Methods). Colours distinguish exons (blue) and promoters (purple). (E) Tumour RNAseq: expression levels of lncRNA genes in different cancer tissues obtained from RNAseq expression data from PCAWG. For B-D, statistical significance was calculated using Wilcoxon test.

First, associations with expected cancer-related features were tested (Figure 4B). CLC genes are significantly more likely to have their transcription start site (TSS) within 100 kb of cancer-associated germline SNPs (“Cancer SNPs 100kb TSS”), and more likely to be either differentially-expressed or epigenetically-silenced in tumours (Yan et al. 2015) (Figure 4B). Intriguingly, we observed a tendency for CLC lncRNAs to be more likely to lie within 1 kb of known cancer protein-coding genes (“CGC 1kb TSS”) – this is explored in more detail below. Furthermore, we found that CLC genes are also significantly closer to non-cancer, phenotype-associated germline SNPs (“NonCancer SNPs 100kb TSS”) in comparison to nonCLC genes (Figure 4B), supporting the biological functionality of CLC genes.

We next investigated the properties of the genes themselves. As seen in Figure 4C, and consistent with our previous findings (Lanzós et al. 2017), CLC genes (“Gene length”) and their spliced products (“Exonic length”) are significantly longer than average. No difference was observed in the ratio of exonic to total length (“Exonic content”), nor overall exon repetitive sequence coverage (“Repeats coverage”), nor GC content.

CLC genes also tend to have greater evidence of function, as inferred from evolutionary conservation. Base-level conservation at various evolutionary depths was calculated for lncRNA exons and promoters (Figure 4D). Across all measures tested, using either average base-level scores or percent coverage by conserved elements, we found that CLC genes’ exons are significantly more conserved than other lncRNAs (Figure 4D). The same was observed for conservation of promoter regions.

High levels of gene expression in normal tissues are known to correlate with lncRNA conservation, and are hypothesized to be a reflection of functionality (Managadze et al. 2011). Additionally, genes with oncogenic roles tend to be highly expressed in cancer samples (Furney et al. 2006). We found that CLC has consistently higher steady-state expression levels across PCAWG tumours (Figure 4E), as well as healthy organs and cultured cell lines (Supplementary Figure 2).

Finally, we investigated whether CLC transcripts might be initiated by any types of Transposable Elements (TEs) (see Materials and methods). We found that CLC TSSs are enriched for one category, “Simple repeats” (Supplementary Figure 3).

### Evidence for genomic clustering of non-coding and protein-coding cancer genes

In light of recent evidence for colocalisation and coexpression of disease-related lncRNAs and protein-coding genes (Tan et al. 2017), we were curious whether such an effect holds for cancer-related lncRNAs and protein-coding genes. We asked, more specifically, whether CLC genes tend to be closer to CGC genes than expected by chance, and whether this is manifested in a more co-regulated expression.

To this aim, we computed TSS-TSS distances from lncRNAs to protein-coding genes and we found that CLC genes on average tend to lie moderately closer to protein-coding genes of all types, compared to nonCLC lncRNAs (Supplementary Figure 4A, B). Since CLC genes are enriched for functional features (i.e. expression and conservation), we couldn’t rule out the possibility that proximity to protein-coding genes is a feature of functional lncRNAs rather than cancer lncRNA genes. In order to further investigate this possibility, we repeated the analysis dividing the nonCLC set into potentially functional nonCLC genes (PF-nonCLC) (nonCLC genes sampled to match CLC expression and conservation, N=149, Supplementary Figure 5) and “other nonCLC” (the rest of nonCLC). Interestingly, when comparing distances to any type of protein-coding genes, both CLC and PF-nonCLC are significantly closer than the rest of lncRNA (Wilcoxon test, P=0.03, 0.007, respectively), being the PF-nonCLC genes the closest ones (median 21.9 kb, 29 kb and 37.8 kb, for PF-nonCLC, CLC, and other nonCLC, respectively) (Supplementary Figure 4C). However, when assessing specifically for distance to CGC genes, only CLC set is significantly closer than the rest of lncRNAs (Wilcoxon test, P=0.0008) and it represents the group with the lowest distance (median 1,122 kb, 1,330kb and 1,607 kb for CLC, PF-nonCLC, and other nonCLC, respectively) (Figure 5A). Thus, although proximity to protein-coding genes seems to be a feature of potentially functional lncRNAs, CLC genes are closer to cancer genes compared to other lncRNAs with similar function-like properties.

**Figure 5:**
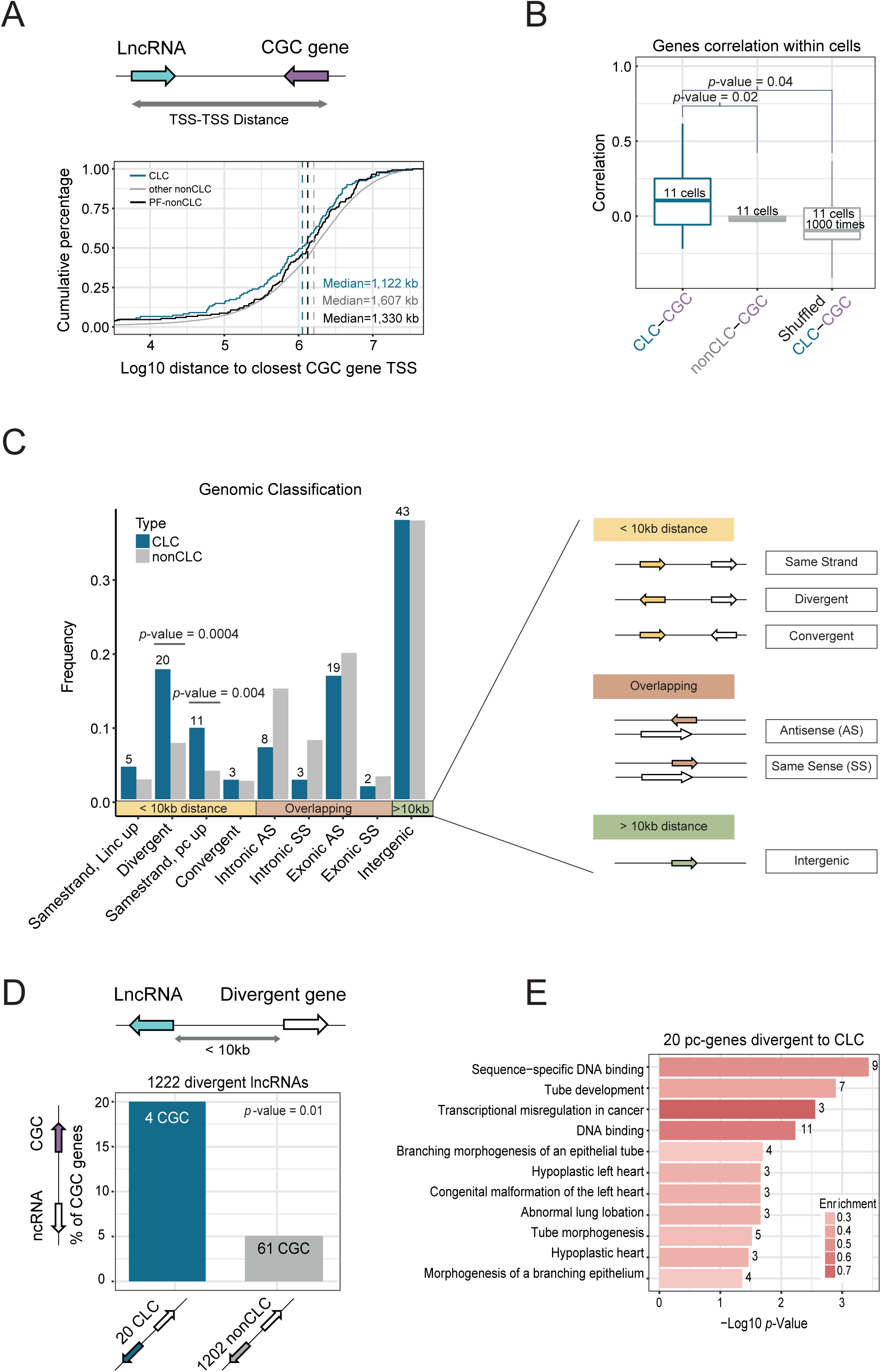
Evidence for genomic clustering of non-coding and protein-coding cancer genes. (A) Cumulative distribution of the genomic distance of lncRNA transcription start site (TSS) to the closest Cancer Gene Census (CGC) (protein-coding) gene TSS. LncRNAs are divided into CLC (n=122), potentially functional nonCLC genes (PF-nonCLC) (n=149), and other nonCLC genes (n=15678) (B) Boxplot shows the distribution of the gene expression correlation between CLC and their closest CGC genes in 11 human cell lines, including two control analyses (distance-matched nonCLC-CGC pairs, and shuffled CLC-CGC pairs). Correlation was calculated for gene pairs within each cell type, using Pearson method. *p*-value for Kolmogorov-Smirnov test is shown. (C) Genomic classification of lncRNAs. Genes are classified according to distance and orientation to the closest protein-coding gene, and these are grouped into three categories: genes closer than 10kb to closest protein-coding gene, genes overlapping a protein-coding gene and intergenic genes (>10kb from closest protein-coding gene). *p*-values for Fisher’s exact tests are shown. (D) The percentage of divergent CLC (left bar) and nonCLC (right bar) genes divergent to a cancer protein-coding gene (CGC). Numbers represent numbers of genes with which the percentage is calculated. *p*-value for Fisher’s exact test is shown. (E) Functional annotations of the 20 protein-coding genes divergent to CLC genes from Panel C. Bars indicate the –log10 (corrected) *p*-value (see Materials and Methods) and are coloured based on the “enrichment”: the number of genes that contain the functional term divided by the total number of queried genes. Numbers at the end of the bars correspond to the number of genes that fall into the category.

It has been widely proposed that proximal lncRNA / protein-coding gene pairs are involved in *cis*-regulatory relationships, which is reflected in expression correlation (Ponjavic et al. 2009). We next asked whether proximal CLC-CGC pairs exhibit this behaviour. An important potential confounding factor, is the known positive correlation between nearby gene pairs (Marques et al. 2013), and this must be controlled for. Using gene expression data across 11 human cell lines, we observed a positive correlation between CLC-CGC gene pairs for each cell type (Figure 5B). To control for the effect of proximity on correlation, we next randomly sampled a similar number of non-CLC lncRNAs with matched distances (TSS-TSS) from the same CGC genes, and found that this correlation was lost (Figure 5B, “nonCLC-CGC”). To further control for a possible correlation arising from the simple fact that both CGC and CLC genes are involved in cancer, and CLC genes are in general enriched for conservation and expression, we next randomly shuffled the CLC-CGC pairs 1000 times, again observing no correlation (Figure 5B, “Shuffled CLC-CGC”). Together these results show that genomically-proximal protein-coding/non-coding gene pairs exhibit an expression correlation that exceeds that expected by chance, even when controlling for genomic distance.

These results prompted us to further explore the genomic localization of CLC genes relative to their proximal protein-coding gene and the nature of their neighbouring genes. Next, we observed an unexpected difference in the genomic organisation of CLC genes: when classified by orientation with respect to nearest protein-coding gene (Derrien et al. 2012), we found a significant enrichment of CLC genes immediately downstream and on the same strand as protein-coding genes (“Samestrand, pc up”, Figure 5C). Moreover, CLC genes are approximately twice as likely to lie in an upstream, divergent orientation to a protein-coding gene (“Divergent”, Figure 5C). Of these CLC genes, 20% are divergent to a CGC gene, compared to 5% for nonCLC genes (P=0.018, Fisher’s exact test) (Figure 5D), and several are divergent to protein-coding genes that have also been linked or defined to be involved in cancer, despite not being classified as CGCs (Supplementary Table 2).

Given this noteworthy enrichment of CGC genes in a divergent configuration to protein-coding genes, we next inspected the latters’ function annotation. Examining their Gene Ontology (GO) terms, molecular pathways and other gene function related terms, we found this group of genes to be enriched in GO terms for “sequence-specific DNA binding”, “DNA binding”, “tube development” and “transcriptional misregulation in cancer” (Figure 5E). These results were confirmed by another, independent GO-analysis suite (see Materials and Methods). Interestingly, three out of the top four functional groups were observed previously in a study of protein-coding genes divergent to long upstream antisense transcripts in primary mouse tissues (Lepoivre et al. 2013).

Thus, CLC genes appear to be non-randomly distributed with respect to protein-coding genes, and particularly their CGC subset.

### Evidence for anciently conserved cancer roles of lncRNAs

In mouse, numerous studies have employed unbiased forward genetic screens to identify genes that either inhibit or promote tumorigenesis (Copeland & Jenkins 2010). These studies use engineered, randomly-integrating transposons carrying bidirectional polyadenylation sites as well as strong promoters. Insertions, or clusters of insertions, called “common insertion sites” (CIS) that are identified in sequenced tumour DNA, implicate the overlapping or neighbouring gene locus as either an oncogene or tumour-suppressor gene. Although these studies have traditionally been focused on identifying protein-coding genes, they can in principle also identify non-coding RNA driver loci.

We thus reasoned that comparison of mouse CISs to orthologous human regions could yield independent evidence for the functionality of human cancer lncRNAs (Figure 6A). To test this, we collected a comprehensive set of CISs in mouse (Abbott et al. 2015), consisting of 2,906 loci from 7 distinct cancer types (Supplementary Table 4). These sites were then mapped to orthologous regions in the human genome, resulting in 1,309 human CISs, or hCISs. 7.3% of these CISs lie outside of protein-coding gene boundaries, and were used for the following analyses.

**Figure 6:**
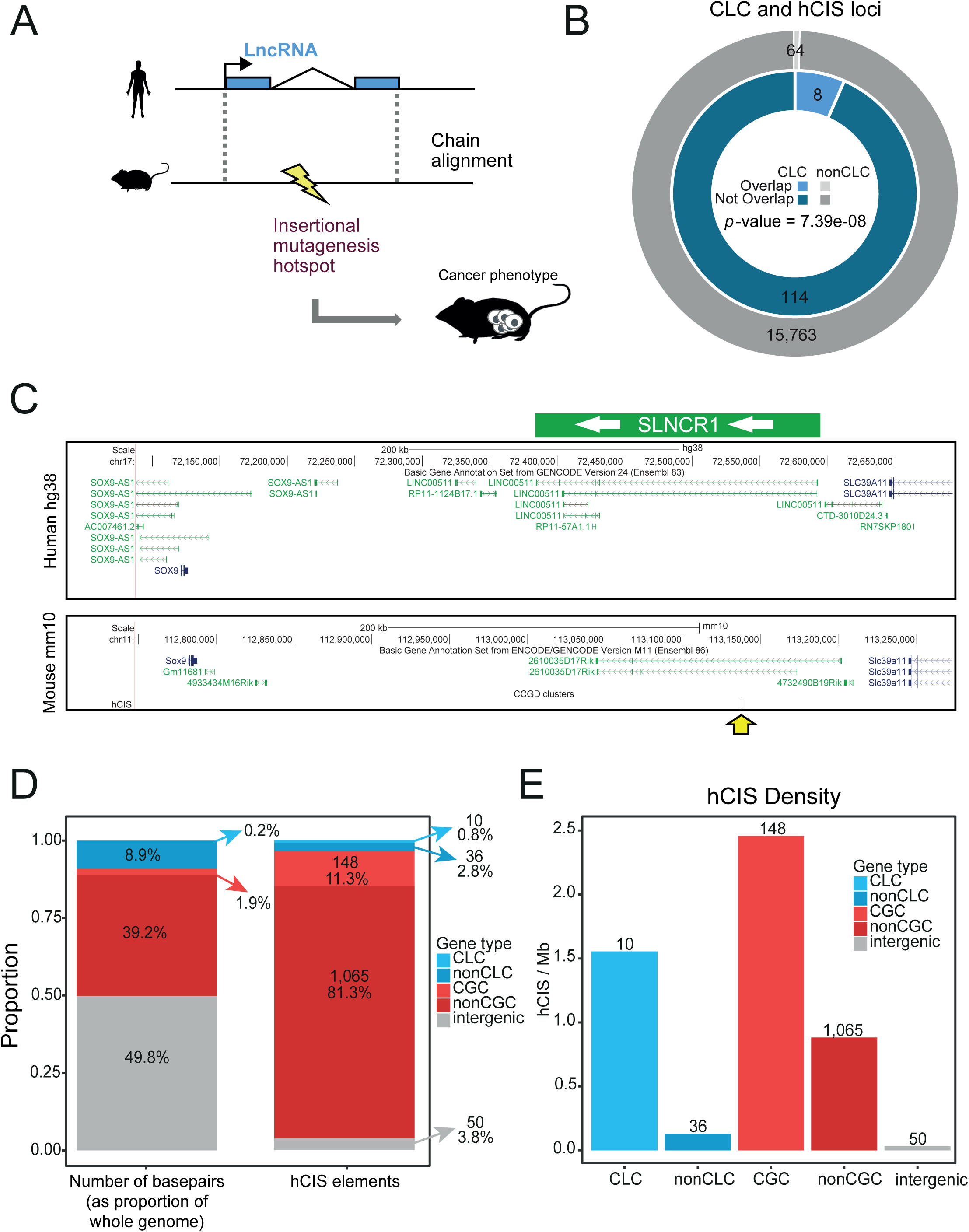
Evidence for ancient conserved cancer roles of lncRNAs. (A) Functional conservation of human CLC genes was inferred by the presence of Common Insertion Sites (CIS), identified in transposon mutagenesis screens, at orthologous regions in the mouse genome. Orthology was inferred from Chain alignments and identified using LiftOver utility. (B) Number of CLC and nonCLC genes that contain human orthologous common insertion sites (hCIS) (see Table 1). Significance was calculated using Fisher’s exact test. (C) UCSC browser screenshot of a CLC gene (*SLNCR1*, ENSG00000227036) intersecting a CIS (yellow arrow). (D) Number of basepairs and number of overlapping hCIS for cancer driver protein-coding genes (CGC), non cancer driver protein-coding genes (nonCGC), cancer related lncRNAs (CLC), rest of GENCODE lncRNAS (nonCLC) and the rest of the genome that do not overlap any of the previous element types (intergenic). Arrows indicate the number of hCIS and the percentage for each element type. (E) Number of overlapping hCIS per megabase of genomic span for each gene class.

Mapping hCISs to lncRNA annotations, we discovered altogether eight CLC genes (6.6%) carrying at least one insertion within their gene span: *DLEU2*, *GAS5*, *MONC*, *NEAT1*, *PINT*, *PVT1*, *SLNCR1*, *XIST* (Table 1). In contrast, just 61 (0.4%) nonCLC genes contained hCISs (Figure 6B). A good example is *SLINCR1*, shown in Figure 6C, which drives invasiveness of human melanoma cells (Schmidt et al. 2016), and whose mouse orthologue contains a CIS discovered in pancreatic cancer. We examined the possibility that hCIS insertions in these CLC genes could in fact be caused by nearby, protein-coding cancer genes. However, none of these eight CLC genes are within 100 kb of a CGC gene, with the exception of *CCAT1* lncRNA, lying 58 kb from *c-MYC* oncogene.

This analysis would suggest that CLC genes are enriched for hCISs; however, there remains the possibility that this is confounded by their greater length. To account for this, we performed two separate validations. First, sets of nonCLC genes with CLC-matched length were randomly sampled, and the number of intersecting hCISs per unit gene length (Mb) was counted (Supplementary Figure 6A). Second, CLC genes were randomly relocated in the genome, and the number of genes intersecting at least one hCIS was counted (Supplementary Figure 6B). Both analyses showed that the number of intersecting hCISs per Mb of CLC gene span is far greater than expected. In contrast, nonCLC genes show a depletion for hCIS sites (Supplementary Figure 6C).

We further compared the enrichment of hCIS in protein-coding genes, lncRNA genes and other intergenic space. Compared to the genomic space they occupy, there is a clear enrichment of hCIS elements in both protein-coding CGC genes, as well as CLC lncRNAs (Figure 6D). Expressed as insertion rate per megabase of gene span, it is clear that CLC genes are targeted more frequently than background intergenic DNA and non-cancer-related protein-coding genes. Of note are the non-background insertion rates for non-cancer-related protein-coding and lncRNA genes, suggesting that there remain substantial numbers of undiscovered cancer genes in both groups.

Together these analyses demonstrate that CLC genes are orthologous to mouse cancer-causing genomic loci at a rate greater than expected by random chance. These identified cases, and possibly other CLC genes, display cancer functions that have been conserved over tens of millions of years since human-rodent divergence.

## Discussion

We have presented the Cancer LncRNA Census, the first controlled set of GENCODE-annotated lncRNAs with demonstrated roles in tumorigenesis or cancer phenotypes.

The present state of knowledge of lncRNAs in cancer, and indeed lncRNAs generally, remains highly incomplete. Consequently, our aim was to create a geneset with the greatest possible confidence, by eliminating the relatively large number of published “cancer lncRNAs” with as-yet unproven causative roles in disease processes. Thus, we used a rather strict definition of cancer lncRNA, being those having direct experimental or genetic evidence for a causative role in cancer phenotypes. By this measure, gene expression changes alone do not suffice. By introducing these well-defined inclusion criteria, we hope to ensure that CLC contains the highest possible proportion of *bona fide* cancer genes, giving it maximum utility for *de novo* predictor benchmarking. In addition, its basis in GENCODE ensures portability across datasets and projects. Inevitably some well-known lncRNAs did not meet these criteria (including *SRA1*, *CONCR, KCNQ1OT1*) (Marchese et al. 2016; Lanz et al. 1999; Higashimoto et al. 2006); these may be included in future when more validation data becomes available. We believe that CLC will complement the established lncRNA databases such as *lncRNAdb*, *LncRNADisease* and *Lnc2Cancer*, which are more comprehensive, but are likely to have a higher false-positive rate due to their more relaxed inclusion criteria (Chen et al. 2013; Quek et al. 2015; Ning et al. 2016).

*De novo* lncRNA driver gene discovery is likely to become increasingly important as the number of sequenced tumours grow. The creation and refinement of statistical methods for driver gene discovery will depend on the available of high-quality true positive genesets such as CLC. It will be important to continue to maintain and improve the CLC in step with anticipated growth in publications on validated cancer lncRNAs. Very recently, CRISPR-based screens (Zhu et al. 2016; Liu et al. 2017) have catalogued large numbers of lncRNAs contributing to proliferation in cancer cell lines, which will be incorporated in future versions.

We used CLC to estimate the performance of *de novo* driver lncRNA predictions from the PCAWG project, made using the ExInAtor pipeline (Lanzós et al. 2017). Supporting the usefulness of this approach, we found an enrichment for CLC genes amongst the top-ranked driver predictions. Extending this to the full set of PCAWG driver predictors, approximately ten percent of CLC genes (9.8%) are called as drivers by at least one method (PCAWG Consortium, Manuscript In Preparation), which is lower to the rate of CGC genes identified (25.1%).

The low rate of concordance between *de novo* predictions and CLC genes may be due to technical or biological factors. Indeed, it is important to state that we do not yet know whether CLC holds “cancer driver” lncRNAs, and indeed, how many such genes exist. In principle, lncRNAs may play two distinct roles in cancer: first, as driver genes, defined as those whose mutations are early and positively-selected events in tumorigenesis; or second, as “downstream genes”, which do make a genuine contribution to cancer phenotypes, but through non-genetic alterations in cellular networks resulting from changes in expression, localisation or molecular interactions. These downstream genes may not display positively-selected mutational patterns, but would be expected to display cancer-specific alterations in expression. A key question for the future is how lncRNAs break down between these two categories, and the utility of CLC in benchmarking *de novo* driver predictions will depend on this. However, the identification of lncRNAs whose silencing or overexpression is sufficient for tumour formation in mouse, would seem to suggest that they are true “driver genes”.

Analysis of the CLC geneset has broadened our understanding of the unique features of cancer lncRNAs, and generally supports the notion that lncRNAs have intrinsic biological functionality. Cancer lncRNAs are distinguished by a series of features that are consistent with both (a) roles in cancer (eg tumour expression changes), and (b) general biological functionality (eg high expression, evolutionary conservation). Elevated evolutionary conservation in the exons of CLC genes would appear to support their functionality as a mature RNA transcript, in contrast to the act of their transcription alone (Latos et al. 2012). Another intriguing observation has been the colocalisation of cancer lncRNAs with known protein-coding cancer genes: these are genomically proximal and exhibit elevated expression correlation. This points to a regulatory link between cancer lncRNAs and protein-coding genes, perhaps through chromatin looping, as described in previous reports for *CCAT1* and *MYC*, for example (Xiang et al. 2014).

One important caveat for all features discussed here is ascertainment bias: almost all lncRNAs discussed have been curated from published, single-gene studies. It is entirely possible that selection of genes for initial studies was highly non-random, and influenced by a number of factors – including high expression, evolutionary conservation and proximity to known cancer genes – that could bias our inference of lncRNA features. This may be the explanation for the observed excess of cancer lncRNAs in divergent configuration to protein-coding genes. However, the general validity of some of the CLC-specific features described here – including high expression and evolutionary conservation – were also observed recent unbiased genome-wide screens (Lanzós et al. 2017; Liu et al. 2017), suggesting that they are genuine.

Despite the relatively low concordance of CLC genes with PCAWG driver predictions, the results of this study strongly support the value and key cancer role of identified lncRNAs in cancer. Most notably, the existence of a core set of eight lncRNAs with independently-identified mouse orthologues with similar cancer functions, is a powerful evidence that these genes are *bona fide* cancer genes, whose overexpression or silencing can drive tumour formation. To our knowledge this is the most direct demonstration to date of anciently-conserved functions and disease roles for lncRNAs. It will be intriguing to investigate in future whether more human-mouse orthologous lncRNAs have been identified in such screens.

## Materials and Methods

### Manual Curation

All lncRNAs in lncRNAdb and those listed in Schmitt and Chang’s recent review article were collected (Quek et al. 2015; Schmitt et al. 2016). To these were added all cases from *LncRNADisease* and *Lnc2Cancer* databases (Chen et al. 2013; Ning et al. 2016). This primary list formed the basis for a manual literature search: all available publications for each gene were identified by keyword search in Pubmed. If publications were found conforming to at least one of the inclusion criteria (below) and the gene has a GENCODE ID, then it was added to CLC, with appropriate information on the associated cancer, biological activity. For the numerous cases where no GENCODE ID was supplied in the original publication, any available ID, or primer or siRNA sequence was used to identify the gene using the UCSC Genome Browser Blat tool (Kent et al. 2002).

Inclusion criteria sufficient to define a cancer lncRNA and link it to a cancer type were:

1. Class t: *In vitro* demonstration that their knockdown and/or overexpression in cultured cancer cells results in changes to cancer-associated phenotypes. These typically include proliferation rates, migration, sensitivity to apoptosis, or anchorage-independent growth.
2. Class v: *In vivo* demonstration that their knockdown and/or overexpression in cancer cells alters their tumorigenicity when injected into animal models.
3. Class g: Germline mutations or variants that predispose humans to cancer.
4. Class s: Somatic mutations that show evidence for positive selection during tumour formation. An additional criteria was allowed to link an lncRNA to a cancer type, only if at least one of the above criteria was already met for another cancer:
5. Class p: Prognosis: The lncRNAs expression is statistically linked to disease progression or response to treatment.

If an lncRNA was found to promote tumorigenesis or cancer phenotype, it was defined as “oncogene” (og). Conversely those found to inhibit such phenotypes were defined as “tumour suppressor” (tsg). Several lncRNAs were found to have both activities recorded in different cancer types, and were given both labels (og/tsg). For every lncRNA-cancer association, a single representative publication is recorded. Finally, it is important to note that no lncRNAs were included based on evidence from previous driver gene discovery studies of the types represented by OncodriveFML, ExInAtor, ncdDetect or others described in PCAWG (Mularoni et al. 2016; Lanzós et al. 2017; Juul et al. 2017) (PCAWG Consortium, Manuscript In Preparation).

CLC set at this stage relies on GENCODE v24 annotation, and therefore all CLC genes have a GENCODE v24 ID assigned. However, data relative to GENCODE v24 was not available for all types of data and analysis used in this study (ie all data relative to PCAWG is based on GENCODE v19). Thus, for some analysis only genes also present in GENCODE v19 could be used (specified in the corresponding methods section) and the total number of genes analysed in these cases is slightly lower (107 instead of 122 CLC genes and 13,503 instead of 15,827 nonCLC).

### LncRNA and protein-coding driver prediction analysis

LncRNA and protein-coding predictions for ExInAtor and the rest of PCAWG methods, as well as the combined list of drivers, were extracted from the consortium database (PCAWG Consortium, Manuscript In Preparation). Parameters and details about each individual methods and the combined list of drivers can be found on the main PCAWG driver publication (PCAWG Consortium, Manuscript In Preparation) and false discovery rate correction was applied on each individual cancer type for each individual method in order to define candidates (*q*-value cutoff of 0.1). This way, we combined the predicted candidates of each individual method in each individual cancer type (including pan-cancer). To calculate sensitivity (percentage of true positives that are predicted as candidates) and precision (percentage of predicted candidates that are true positives) for lncRNA and protein-coding predictions we used the CLC and CGC (COSMIC v78, downloaded Oct, 3, 2016) sets, respectively. To assess the statistical significance of sensitivity rates, we used Fisher’s exact test.

### Feature Identification

We compiled several quantitative and qualitative traits of GENCODE lncRNAs and used them to compare CLC genes to the rest of lncRNAs (referred to as “nonCLC”). Analysis of quantitative traits were performed using Wilcoxon test while qualitative traits were tested using Fisher' exact test. These methods principally refer to Figure 4 and 5 as well as Supplementary Figures 2, 3, 4 and 5.

#### Cancer SNPs

On October, 4, 2016, we collected all 2,192 SNPs related to “cancer”, “tumour” and “tumor” terms in the NHGRI-EBI Catalog of published genome-wide association studies (Hindorff et al. 2009; Welter et al. 2014) (https://www.ebi.ac.uk/gwas/home). Then we calculated the closest SNP to each lncRNA TSS using *closest* function from Bedtools v2.19 (Quinlan & Hall 2010) (GENCODE v24).

#### NonCancer SNPs

On July, 31, 2017, we collected all 29,813 SNPs not related to “cancer”, “tumour” and “tumor” terms in the NHGRI-EBI Catalog of published genome-wide association studies (Hindorff et al. 2009; Welter et al. 2014) (https://www.ebi.ac.uk/gwas/home). Then we calculated the closest SNP to each lncRNA TSS using *closest* function from Bedtools v2.19 (Quinlan & Hall 2010)(GENCODE v24).

#### Epigenetically-silenced lncRNAs

We obtained a published list of 203 cancer-associated epigenetically-silenced lncRNA genes present in GENCODE v24 (Yan et al. 2015). These candidates were identified due to DNA methylation alterations in their promoter regions affecting their expression in several cancer types.

#### Differentially expressed in cancer

We collected a list of 3,533 differentially-expressed lncRNAs in cancer compared to normal samples (Yan et al. 2015) (GENCODE v24).

#### Sequence / gene properties

Exonic positions of each gene were defined as the projection of the union of exons from all its transcripts. Introns were defined as all remaining non-exonic nucleotides within the gene span. Repeats coverage refers to the percent of exonic nucleotides of a given gene overlapping repeats and low complexity DNA sequence regions obtained from RepeatMasker data housed in the UCSC Genome Browser (Tyner et al. 2017). Exonic content refers to the fraction of total gene span covered by exons. For this section we used GENCODE v19.

#### Evolutionary conservation

Two types of PhastCons conservation data were used: base-level scores and conserved elements. These data for different multispecies alignments (GRCh38/hg38) were downloaded from UCSC genome browser (Tyner et al. 2017). Mean scores and percent overlap by elements were calculated for exons and promoter regions (GENCODE v24). Promoters were defined as the 200nt region centred on the annotated gene start.

#### Expression

We used polyA+ RNA-seq data from 10 human cell lines produced by ENCODE (Djebali et al. 2012; ENCODE Project Consortium et al. 2012), from various human tissues by the Illumina Human Body Map Project (HBM) (www.illumina.com; ArrayExpress ID: E-MTAB-513), and from cancer samples from PCAWG RNAseq expression data (PCAWG Consortium, Manuscript In Preparation). In this last case, for each cancer type we computed the expression mean of genes across all RNAseq samples belonging to that cancer type (GENCODE v19).

#### Transposable elements

We downloaded 5,520,016 transposable elements from the UCSC table browser (Karolchik et al. 2004) on August, 3, 2017. We separated them by element types and counted how many of them intersected or not with the transcription start sites of CLC and nonCLC genes, in order to detect any association with the Fisher' exact test.

#### Distance to protein-coding genes and CGC genes

For each lncRNA we calculated the TSS to TSS distance to the closest protein-coding gene (GENCODE v24) or CGC gene (downloaded on October, 3, 2016 from Cosmic database) (Futreal et al. 2004) using *closest* function from Bedtools v2.19 (Quinlan & Hall 2010). In order to divide nonCLC genes into potentially functional nonCLC (PF-nonCLC) and others, we sampled the list of all nonCLC genes to get a subsample that has a matched distribution to CLC genes in conservation (% of conserved elements, from Vertebrate Multiz Alignment 100 Species from UCSC genome browser data, in exonic regions). Then we sampled again the resulting subset to get a final subset that also matches CLC genes in terms of expression (median of expression across 16 human tissues, data from Illumina Human Body Map Project (HBM)). To create the nonCLC samples we used the *matchDistribution* script: https://github.com/julienlag/matchDistribution.

#### Coexpression with closest CGC gene

We took CLC-CGC gene pairs whose TSS-TSS distance was <200kb. RNAseq data from 11 human cell lines from ENCODE was used to assess expression levels (Djebali et al. 2012; ENCODE Project Consortium et al. 2012). ENCODE RNAseq data were obtained from ENCODE Data Coordination Centre (DCC) in September 2016, https://www.encodeproject.org/matrix/?type=Experiment. All data is relative to GENCODE v24. We calculated the expression correlation of gene pairs within each of the 11 cell lines, using the Pearson measure. To control for the effect of proximity, we randomly sampled a subset of nonCLC-CGC pairs matching the same TSS-TSS distance distribution as above, and performed the same expression correlation analysis (“nonCLC-CGC”). Finally, to further control for the fact that CLC and CGC are both cancer genes, which may influence their expression correlation, we shuffled CLC-CGC pairs 1000 times, and tested expression correlation for each set (“Shuffled CLC-CGC”).

#### Genomic classification

We used an in-house script to classify lncRNA transcripts into different genomic categories based on their orientation and proximity to the closest protein-coding gene (GENCODE v24): a 10 kb distance was used to distinguish “genic” from “intergenic” lncRNAs. When transcripts belonging to the same gene had different classifications, we used the category represented by the largest number of transcripts.

#### Functional enrichment analysis

The list of protein-coding genes (GENCODE v24) that are divergent and closer than 10 kb to CLC genes (or nonCLC) was used for a functional enrichment analysis (20 unique genes in the case of CLC analysis and 1202 in the case of nonCLC analysis). We show data obtained using g:Profiler web server (Reimand et al. 2016), g:GOSt, with default parameters for functional enrichment analysis of protein-coding genes divergent to CLC and using Bonferroni correction for protein-coding gene divergent to nonCLC. For CLC analysis we performed the same test with independent methods: Metascape (http://metascape.org) (Tripathi et al. 2015) and GeneOntoloy (Panther classification system)(Mi et al. 2013; Mi et al. 2017). In both cases similar results were found.

### Mouse mutagenesis screen analysis

We extracted the genomic coordinates of transposon common insertion sites (CISs) in Mouse (GRCm38/mm10) http://ccgd-starrlab.oit.umn.edu/about.php (Abbott et al. 2015). This database contains target sites identified by transposon-based forward genetic screens in mice. LiftOver (Kent et al. 2002) was used at default settings to obtain aligned human genome coordinates (hCISs) (GRCh38/hg38). We discarded hCIS regions longer than 1000 nucleotides and those that overlap protein-coding genes, and intersected the remainders with the genomic coordinates CLC and nonCLC genes.

To correctly assess the statistical enrichment of CLC in hCIS regions, we performed two control analyses:

#### Randomly repositioning of CLC and nonCLC genes

We randomly relocated CLC/nonCLC genes 10,000 times within the whole genome using the tool *shuffle* from BedTools v19 (Quinlan & Hall 2010). In each iteration, we calculated the number of genes that intersected at least one hCIS, and created the distribution of these simulated values. Finally, we calculated an empirical *p*-value by counting how many of the simulated values were higher or equal than the real values. This analysis was performed separately for CLC and nonCLC genes.

#### Length-matched sampling

To calculate if the enrichment of hCIS intersecting genes in CLC set is higher and statistically different from nonCLC set, while controlling by gene length, we created 1000 samples of nonCLC genes with the same gene length distribution as CLC genes. Each sample was intersected with hCIS, and the number of intersecting hCISs per Mb of gene length was calculated. To create the nonCLC samples we used the *matchDistribution* script: https://github.com/julienlag/matchDistribution.

## Acknowledgements

We wish to thanks Julien Lagarde (CRG) for help and advice in bioinformatic analysis. We acknowledge Romina Garrido (CRG), Deborah Re (DKF), Silvia Roesselet (DKF) and Marianne Zahn (Inselspital) for administrative support. We thank Ivo Buchhalter (DKFZ) and Sandra Koser (DKFZ) for preprocessing the SNV and expression data for the integrated analysis. Iñigo Martincorena (Sanger Institute) kindly provided the script for analysing driver prediction sensitivity. A.L. is supported by pre-doctoral fellowship FPU14/03371. This research was supported by the NCCR RNA & Disease funded by the Swiss National Science Foundation.

## Contributions

RJ conceived the project, performed manual annotation of CLC, and supervised with advice and suggestions of JS-P, LF and CH. JCF and AL performed the feature analysis and evolutionary analysis. AL performed mutation analysis. RJ, AL and JCF drafted the manuscript and prepared the figures and supplementary material. All authors read and approved the final draft.

## Supplementary Figure Legends

**Supplementary Figure 1: CLC summary statistics**. (A) Barplot showing the non-redundant number of genes in CLC broken down by supporting evidence types. p: prognostic; t: in vitro; v: in vivo; g: germline mutations; s: somatic mutations. (B) Similar as previous, but with (redundant) number of genes per individual evidence type. (C) Histogram of genes broken down by their number of associated cancer types. (D) Histogram of cancer types, by their (redundant) number of associated lncRNAs.

**Supplementary Figure 2: CLC genes are highly expressed**. Panels display feature analysis results similar to Figure 4 using other datasets. (A) Panel displaying the log fold difference between CLC and nonCLC genesets (*y*-axis) and statistical significance by Wilcoxon test (*x*-axis) when comparing RNAseq expression levels in human tissues (each dot represents a different tissue) from Human Body Map data. (B) Same than previous panel for expression data in human cell lines instead of tissues, from ENCODE RNAseq data.

**Supplementary Figure 3: CLC TSSs association with Transposable Elements.** Figure shows the comparison of the intersection of each category of Transposable elements with transcription start sites (TSS) of CLC and nonCLC genes. *y*-axis shows the log2 of the Odds Ratio obtained by comparing CLC to nonCLC by Fisher’s exact test; *x-*axis displays the estimated *p*-value from the same test.

**Supplementary Figure 4: CLC lncRNAs tend to be closer to protein-coding genes.** (A) Cumulative distribution of the genomic distance from CLC and nonCLC genes, to the closest protein-coding gene (NB this may be a CGC gene or not). Distances are defined as the distance of the annotated transcription start site (TSS) of each gene in the pair. *p*-value for Wilcoxon test is shown. (B) Same as (A) for genomic distance to closest CGC genes. (C) Same than A dividing nonCLC genes into two groups: potentially functional nonCLC (PF-nonCLC) (those nonCLC genes that are in the same range of expression and conservation than CLC genes) and other nonCLC (the rest of nonCLC genes).

**Supplementary Figure 5: sampling nonCLC genes**. (A) Density plot comparing the percentage of PhastCons conserved elements in lncRNA exons of CLC genes and a subset of nonCLC sampled to match CLC conservation distribution. (B) Same than A but comparing the median of RNAseq expression values across 16 human tissues. NonCLC subset here is sampled from the subset obtained after matching conservation distribution.

**Supplementary Figure 6: hCIS enrichment corrected by gene length.** (A) Distribution of the number of intersecting hCIS per Megabase (Mb) of total gene length, for 1000 subsets of nonCLC genes with same length distribution as CLC genes (grey). Vertical blue line represents the overall value for CLC geneset: 1.42 hCIS sites per Mb of gene span. (B) Distribution of the number of genes overlapping a hCIS after 10,000 genomic randomizations of CLC genes (blue). Vertical black line represents the observed number of CLC genes (8) that intersect a hCIS. (C) Distribution of the number of intersecting genes with a hCIS after 10,000 genomic randomizations of nonCLC genes (grey). Vertical black line represents the observed number of nonCLC genes that intersect a hCIS (64).

## Supplementary Tables

**Supplementary Table 1: full CLC set.**

**Supplementary Table 2: CLC – protein-coding pairs.**

**Supplementary Table 3: GO analysis for protein-coding genes divergent to nonCLC genes.**

**Supplementary Table 4: Counts of mouse CIS per cancer type.**

